# Sexual selection drives the evolution of wing interference patterns

**DOI:** 10.1101/497115

**Authors:** MF Hawkes, E Duffy, R Joag, A Skeats, J Radwan, N Wedell, MD Sharma, DJ Hosken, J Troscianko

## Abstract

The seemingly transparent wings of many insects have recently been found to display dramatic structural coloration. These structural colours (wing interference patterns: WIPs) may be involved in species recognition and mate choice, yet little is known about the evolutionary processes that shape them. Additionally, existing research has been restricted by analysing WIPs without due consideration of how they are actually perceived by the viewers’ colour vision. Here, we use multispectral digital imaging and a model of *Drosophila* vision to compare WIPs of male and female *Drosophila simulans* from replicate populations forced to evolve with or without sexual selection for 68 generations. We show for the first time that WIPs modelled in *Drosophila* vision evolve in response to sexual selection, and confirm that WIPs correlate with male sexual attractiveness. These findings add a new element to the otherwise well described *Drosophila* courtship display and confirm that wing colours evolve through sexual selection.

## Introduction

Animal colour patterns are important sources of information that are used in a range of signalling contexts including species recognition (Barraclough *et al.* 1995), intrasexual competition (Siefferman & Hill 2005), and mate choice (Houde 1997). When colour patterns are subject to sexual selection, colouration covaries with sexual fitness-components, and colour can be part of multi-modal courtship displays (Candolin 2003). Wing interference patterns (WIPs) are a newly discovered visual component of many insect wings that are thought to act as visual displays (Figure 1). They have been recorded in several *Drosophila* species (Shevtsova *et al.* 2011) and possibly represent previously unrecognised sexual signals in otherwise well-described *Drosophila* courtship displays - which also involve species-specific movement, song, olfaction, and taste (Shevtsova *et al.* 2011; Greenspan *et al.* 2000; Katayama *et al.* 2011). WIPs are a form of structural colouration produced by thin-film interference where light striking the wing is refracted and reflected in such a way that the wavelength of the reflected light is dependent on the thickness of the chitinous membrane of the wing (Shevtsova *et al.* 2011). As a result, variation in wing thickness, along with other structural variation including hair placement and venation, determines variation in reflected colour (Shevtsova *et al.* 2011).

**Figure 1;.**
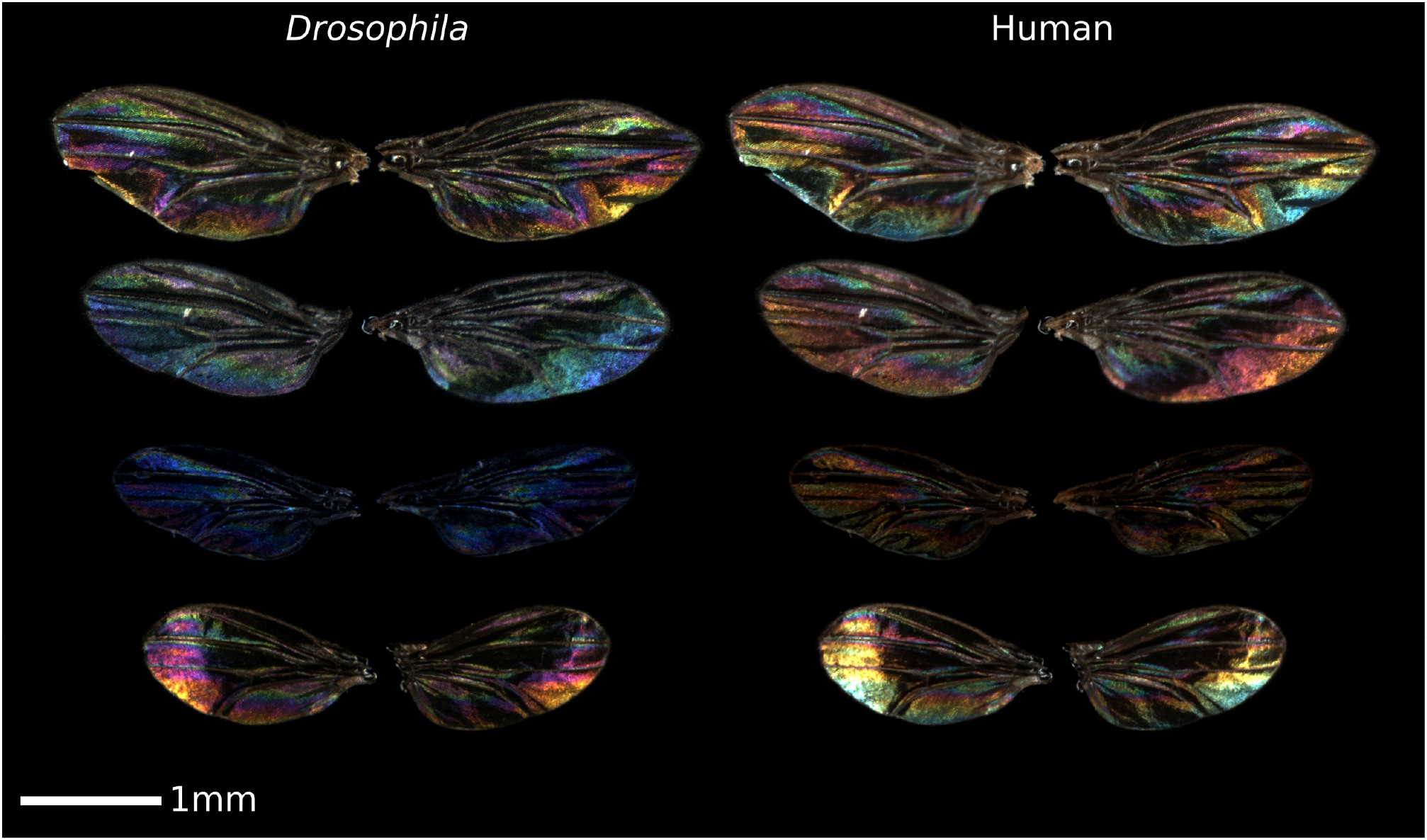
Examples of *Drosophila simulans* wing interference patterns (WIPS) photographed for this study using a customised multispectral photography system. Like many insects, *Drosophila* can see into the ultraviolet range, but not the human ‘red’ range of the spectrum. The images in the left-hand column show ‘false colour’ *Drosophila* vision, where the red, green and blue values correspond to normalised Rh6, Rh5, and Rh4 cone-catch quanta respectively. Images in the right-hand column show the same wings in ‘normal’ human-vision colours.

Recent work shows that WIPs within the human-visible spectrum are heritable and subject to sexual selection via female mate choice in *D. melanogaster* (Katayama *et al.* 2011), but generally little is known about the selective forces that shape WIPs and how they might respond to any such selection. Furthermore, despite evidence that WIPs can be sexual signals, all work to date has used uncalibrated digital images where pixel colour values do not correspond linearly with radiance, making objective colour measurement extremely problematic (Stevens *et al.* 2007). Additionally, no work has yet investigated WIPs explicitly within the spectral sensitivities of the photoreceptors in the *Drosophila* visual system, and so drawing clear biological conclusions about which WIP elements are under selection and how they might evolve is currently not possible.

The *Drosophila* eye contains five main types of photoreceptor, each expressing a single opsin gene; rhodopsins 1 and 3 through 6 (Rh1 and Rh3 through Rh6) (Schnaitmann *et al.* 2013). One is thought to be achromatic with broadband spectral sensitivity to both human-visible and UV light (Rh1), although this may also be used in colour processing (Schnaitmann *et al.* 2013), two have narrow peak sensitivities in the human-visible spectrum roughly corresponding to green (Rh6) and blue (Rh5) light, and two have narrow peak sensitivities in the UV spectrum at shorter (Rh3) and longer (Rh4) wavelengths (Schnaitmann *et al.* 2013, Rister *et al.* 2013). These photoreceptors are arranged into bundles of cells called ommatidia, consisting of a central column of two narrow peak photoreceptors (Rh6 and Rh4, or Rh5 and Rh3) encircled by six Rh1 photoreceptors. Almost all ommatidia are defined as being either ‘pale’ (expressing Rh3 and Rh5) or ‘yellow’ (expressing Rh4 and Rh6) (Rister *et al.* 2013; Hardie 1979; Salcedo *et al.* 1999; Wernet *et al.* 2007). Any investigation of *Drosophila* WIPs needs to take into account these attributes of the visual system if we are to understand any potential signalling roles they might have.

Mating signals and sexual selection has been extensively studied in *D. simulans* (Spieth 1974; Markow *et al.* 1996; Ingleby *et al.* 2013a; Ingleby *et al.* 2013b; Ingleby *et al.* 2013c; Taylor *et al.* 2010; Sharma *et al.* 2012a; Taylor *et al.* 2007; Taylor *et al.* 2008; Sharma *et al.* 2010), but WIPs have not been investigated and incorporated into this framework. Female *D. simulans* are polyandrous, largely determine whether copulation occurs and have a strong preference for certain male genotypes, but do not show clear mate-preference based on male size (Spieth 1974; Ingleby *et al.* 2013b; Ingleby *et al.* 2013c; Taylor *et al.* 2010; Sharma *et al.* 2012a; Taylor *et al.* 2007; Taylor *et al.* 2008; Sharma *et al.* 2010). Here we investigated the impacts of sexual selection on WIPs in *D. simulans.* Calibrated digital imaging with *Drosophila* colour-vision modelling was used to capture WIP colour data as per-pixel ‘cone-catch quanta’ that describe the degree to which the photoreceptors of the *Drosophila* eye are expected to respond to the light (UV and human visible wavelengths) reflected from each wing. WIP colours can vary dramatically within and between wings, however the exact nature of *Drosophila* colour processing is poorly understood. We therefore measured four visual aspects of WIPs likely to be biologically relevant to *Drosophila* visual processing, which were, wing luminance (perceived brightness), luminance contrast, average colour (hue), and colour contrast (variation in hue: see methods for further details). Using male and female wings from experimental populations that had evolved with and without sexual selection (polyandrous and monogamous populations respectively). We provide the first direct evidence that sexual selection can drive the evolution of WIPs within wavelengths of light visible to the *Drosophila* visual system and show that variation in WIPs correlates with male sexual attractiveness.

## Materials & Methods

### Experimental populations

To investigate the ability of sexual selection to drive the evolution of WIPs, we established replicate experimental populations of *D. simulans* that evolved under either enforced monogamy (1♂:1♀, relaxed sexual selection on males) (n=4), or under enforced polyandry (4♂:1♀, elevated sexual selection on males) (n=4) for 68 non-overlapping generations. This is a standard technique for manipulating the opportunity for sexual selection and allows the action of both pre- and post-copulatory selection (Sharma *et al.* 2012b; Holland & Rice 1999; Hosken *et al.* 2001; Crudgington *et al.* 2005; Tilszer *et al.* 2006).

In each generation, males and females were housed in mating vials at their treatment-specific sex ratio for six days (elevated sexual selection: n=60 per replicate. Relaxed sexual selection: n=64 per replicate). More mating vials were included in the relaxed treatment to equalise the effective population size (*N*_e_) between treatments (Sharma *et al.* 2012b). Females were then haphazardly selected to be transferred to treatment- and replicate-specific oviposition vials and housed at a standardised density for 48 hours. Virgin adults were collected from oviposition vials after eclosion under light CO_2_ anaesthesia and separated by sex before being haphazardly assigned to new mating vials for the next generation. Before wings were dissected and photographed all experimental populations were reared for a single generation in mating vials at a standard density (2♂:2♀) to reduce the likelihood of environmental or maternal effects confounding the results (Magalhães *et al.* 2011).

Experimental populations were originally derived from a stock population of *D. simulans* established from flies originally collected in Australia in 2004 after screening with tetracycline to eliminate *Wolbachia* infection. *Wolbachia* infection has been associated with several deleterious effects on fitness in *D. simulans* (Snook *et al.* 2000; de Crespigny & Wedell 2006), and can induce cytoplasmic incompatibility in crosses with differences in infection status or strain (Werren 1997; Werren *et al.* 2008). All flies were housed at a temperature of 25°C under a 12:12hr light:dark cycle on an oatmeal based food media.

We dissected and photographed a total of 480 pairs of wings from 240 individuals. 36 wings were excluded from analyses due to objects obscuring the wing (e.g. fibres) or wing damage. Final sample sizes were: males evolving with sexual selection n=55; males without sexual selection n=58; females with sexual selection n=57; and females without sexual selection n=56 (all groups consisted of individuals sampled from each of 4 replicate populations per treatment).

### Wing interference pattern imaging

Wings were photographed in a custom-built assembly using a calibrated Canon 7D camera that had been converted to full-spectrum sensitivity by replacing the sensor’s visible-band pass filter with a quartz sheet (conversion by Advanced Camera Systems, Norfolk, UK). The camera was fitted with a Novoflex Noflexar 35mm lens that transmits in the visible and ultraviolet (UV) range, reverse-mounted on a helicoid to achieve a suitable magnification. Photographs were taken through a Baader UV/IR cut filter that transmits in the human visible range (400-700 nm), and then through a Baader Venus-U filter that only transmits in the UV (UV, 310-390 nm) range.

Wing interference patterns change dramatically as the angle of the wing, light source and viewing angle change under direct (e.g. point source) illumination. We therefore used a custom-built lighting system that provided uniform, diffuse lighting to create standardised illumination and viewing conditions. The lighting assembly used an Iwasaki eyeColor metal halide arc lamp modified to emit UV light by removal of its UV/IR filter. This bulb is designed to match the Commission on Illumination (CIE) standard D65 illuminant, so recreates natural illumination. The bulb was positioned inside a stainless-steel spherical reflector directly below the sample that focussed light onto a ring of raw white polytetrafluoroethylene plastic sheet around the lens, simulating a ring-flash. Critically, this light source created standardised and uniformly diffuse illumination that matches natural conditions. The dorsal surfaces of wings were photographed in pairs on a dark, spectrally flat polymethyl methacrylate background that contained a scale-bar.

### Image processing

Most imaging systems create photographs for viewing on non-linear, low dynamic range displays using 8-bits per channel colour spaces. However, such images are also non-linear, meaning the pixel values do not correspond linearly with radiance, which in turn makes them unsuitable for objective colour measurement (Stevens *et al.* 2007). Standard Red-Green-Blue (RGB) systems are also unsuitable for modelling *Drosophila* vision because they do not capture the UV portion of the spectrum to which *Drosophila* are sensitive, and previous analyses have included the red portion of the spectrum, which the flies are unable to detect (Briscoe & Chittka 2001). We therefore processed our whole-wing images using our Multispectral Image Analysis and Calibration Toolbox for ImageJ (Schneider *et al.* 2012), which enables image calibration, first controlling for lighting conditions and then converting images to animal cone-catch quanta (Troscianko & Stevens 2015).

We used the toolbox to combine the visible and ultraviolet whole-wing images into aligned, normalised multispectral stacks, and then used a cone-mapping approach to convert these images to *“Drosophila* vision” (i.e. *Drosophila* cone-catch quanta). Images were normalised (i.e. converted to relative reflectance images that control for lighting conditions) by measuring the background grey in each image, which was in turn calibrated against a Spectralon 99% reflectance standard (Labsphere). Briefly, the cone-mapping process uses the known spectral sensitivities of the camera to estimate the camera’s response to a database of thousands of natural reflectance spectra illuminated using the CIE standard D65 illuminant following the von Kries correction. In addition, the *Drosophila* cone-catch quanta were calculated for the same illuminant using *Drosophila* spectral sensitivities (Schnaitmann *et al.* 2013; Salcedo *et al.* 1999). A polynomial model was then fitted between camera and *Drosophila* vision. The model reported R^2^ values >0.993 for all five receptor classes. For more information on the methodology see Troscianko & Stevens (2015).

Mean luminance was calculated as the mean Rh1 cone-catch quanta pixel estimates for each wing, and luminance contrast was the standard deviation in these estimates. Hue was based on four empirically validated opponent colour channels (Rh5-Rh3, Rh6-Rh4, Rh6-Rh1, and Rh1-Rh4) (Schnaitmann *et al.* 2013). Cone-catch images were converted to each of these four opponent channel images (e.g. the green-blue opponent channel was calculated at each pixel as: Rh6/(Rh6+Rh5)) (Spieth 1974). The mean and standard deviation in opponent channel pixel values across each wing were then used to analyse wing colour and colour contrast respectively using principal component analysis (see below). Wings of a single colour would therefore have a high average colour, but low colour contrast, while wings containing multiple colours would have a high colour contrast. NB the variance analyses were comparing mean variances across treatments rather than strictly comparing the variances within treatments (see Hosken *et al.* 2018 for the importance of this distinction).

### Attractiveness assay

After 55 generations of experimental evolution (13 generations before wings were measured – tests were staggered for logistic reasons), virgin males from each experimental population were collected and housed alone until sexually mature. Attractiveness was then measured using standard protocols (Ingleby *et al.* 2013b; Taylor *et al.* 2010; Sharma *et al.* 2012a; Taylor *et al.* 2007) with virgin females from stock populations used as testers (i.e. females that had not been subjected to experimental evolution). In brief, females should mate with attractive males more quickly and we used mating latency (time from pairing until mating: log transformed) as our measure of male attractiveness (Ingleby *et al.* 2013b; Taylor *et al.* 2010; Sharma *et al.* 2012a; Taylor *et al.* 2007; Hosken *et al.* 2008; Kyriacou & Hall 1986; Barth *et al.* 1997; Ritchie *et al.* 1999). A mean latency per population was calculated and then populations were ranked on these means and the rank sum for populations evolving with and with-out sexual selection were then compared using a Mann-Whitney rank-sum test.

### Statistical analyses

All statistical analyses were performed in Statview/SuperAnova (attractiveness) or R version 3.1.2 (R Core Team 2012), where General Linear Mixed Models (GLMM) were implemented in the package *lme4* (Bates *et al.* 2015).

Wing luminance, luminance contrast, colour principal components and colour contrast principal components were compared between sexes and treatments with GLMMs fitted with sex, treatment, and their interaction as fixed effects, and population replicate and individual fly ID as random effects. Fixed effects were tested for significance using the ANOVA function in the *car* package (Fox & Weisberg 2011). Where a significant sex by treatment interaction was present, Tukey contrasts adjusted for multiple comparisons were obtained from the GLMMs using the *lsmeans* package (Lenth 2016). Where no significant interaction between sex and treatment was found, significant GLMM terms explaining differences in WIP traits are reported. Luminance values are presented as Rh1 cone-catch quanta from zero to one. Full GLMM model tables are available in Tables S1-10.

Principal Component Analyses (PCA) were conducted on opponent channel data to reduce the dimensionality of the dataset and account for high levels of correlation between the cone-catch values across opponent channels. Principal components (PCs) derived from the PCA were considered biologically significant if their associated eigenvalue was greater than 1.0 (Norman & Streiner 2008), and the loading of PCs was considered significant if greater than 0.35 (Tabachnick & Fidell 2013). Statistical testing of the principal component data was conducted in the same manner as for the cone-catch quanta data.

## Results

The effect of sexual selection on the brightness of wings (luminance measured by the mean cone-catch values of the broadband photoreceptor Rh1) was dependent on sex (i.e. there was a sex-by-treatment interaction: GLMM, *χ*^2^_1_=9.63, *p*=0.002). The WIPs of males evolving with sexual selection had significantly higher mean luminance values than the WIPs of males evolving without sexual selection (LSMeans, *t* ratio=3.93, p<0.001; Figure 2). In contrast, there was no difference in mean luminance between the WIPs of females evolving with or without sexual selection (LSMeans, *t* ratio=0.44, p=0.97). The luminance of females WIPs were also similar to males evolving without sexual selection (LSMeans, *t* ratio<1.38, p>0.51), but females tended to differ from sexual selection males (LSMeans, *t* ratio=-2.46, p=0.047 and *t* ratio=-2.25, p=0.12 for sexual and non-sexual selection females respectively) (full statistical models for this and subsequent analyses are presented in the supplementary data).

**Figure 2.**
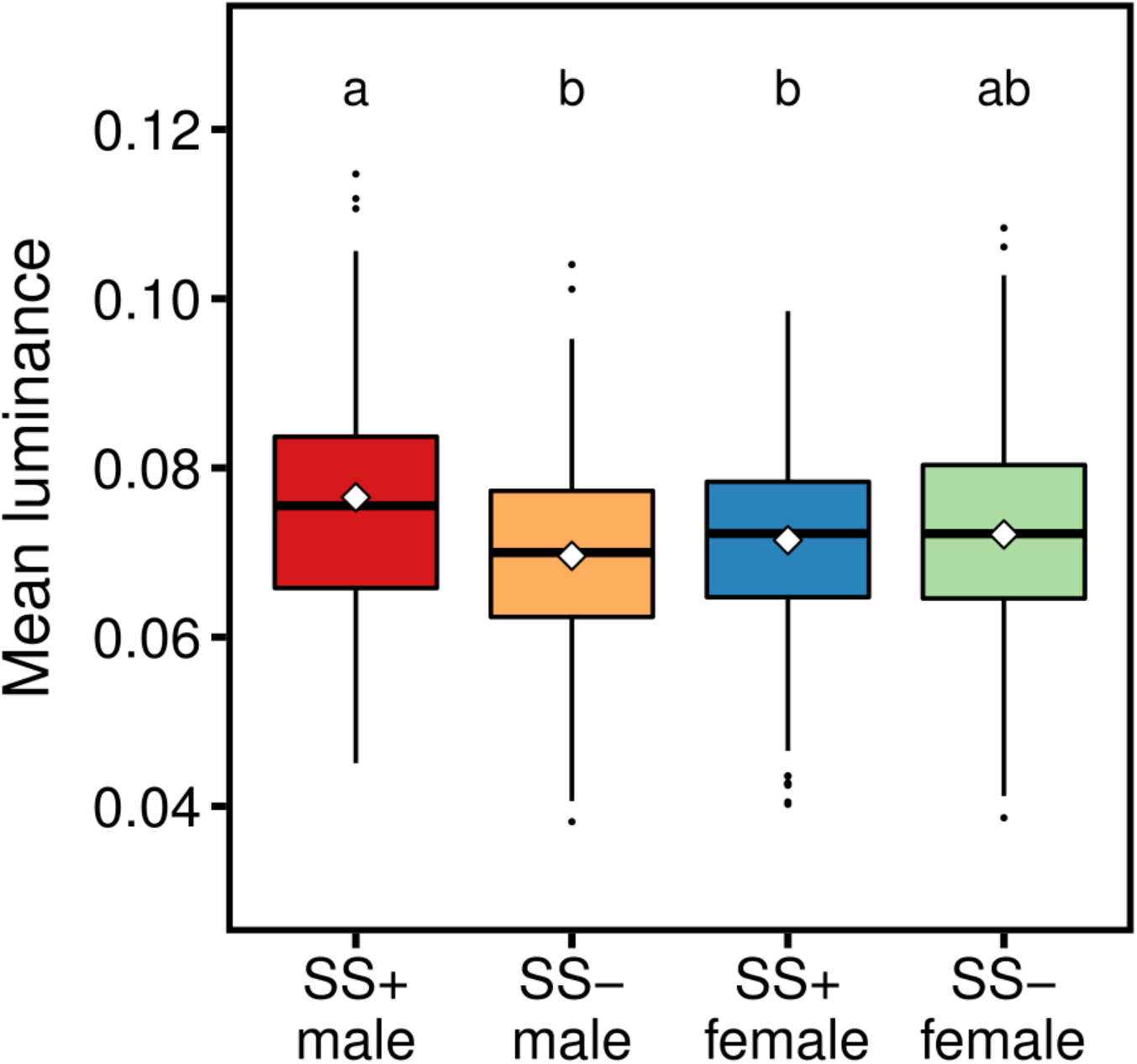
Mean luminance of WIPs as measured by average stimulation of the broadband Rh1 photoreceptor in the *Drosophila* visual system. Boxes represent the interquartile range, black bars are medians, white diamonds are means. SS+ = flies from populations evolving with sexual selection, SS- = flies from populations evolving without sexual selection. Differences in letter annotation denote significance at *p*<0.05.

The effect of sexual selection on the brightness contrast (luminance contrast measured by the standard deviation of the cone-catch values of the broadband receptor Rh1) also showed a sex by treatment interaction (GLMM, *χ*^2^_1_=6.84, *p*=0.009). The WIPs of males evolving with sexual selection had significantly higher luminance contrast than those of males evolving without sexual selection (LSMeans, *t* ratio=4.69, p<0.001; Figure 3), but there was no difference between the WIPs of females evolving with or without sexual selection (LSMeans, *t* ratio=1.02, *p*=0.74; Figure 2). The brightness contrast of females WIPs were again similar to males evolving without sexual selection (LSMeans, *t* ratio<1.86, p>0.25), and again females tended to differ from sexual selection males (LSMeans, *t* ratio=-2.46, p=0.073 and *t* ratio=-3.38, p=0.005 for sexual and non-sexual selection females respectively).

**Figure 3.**
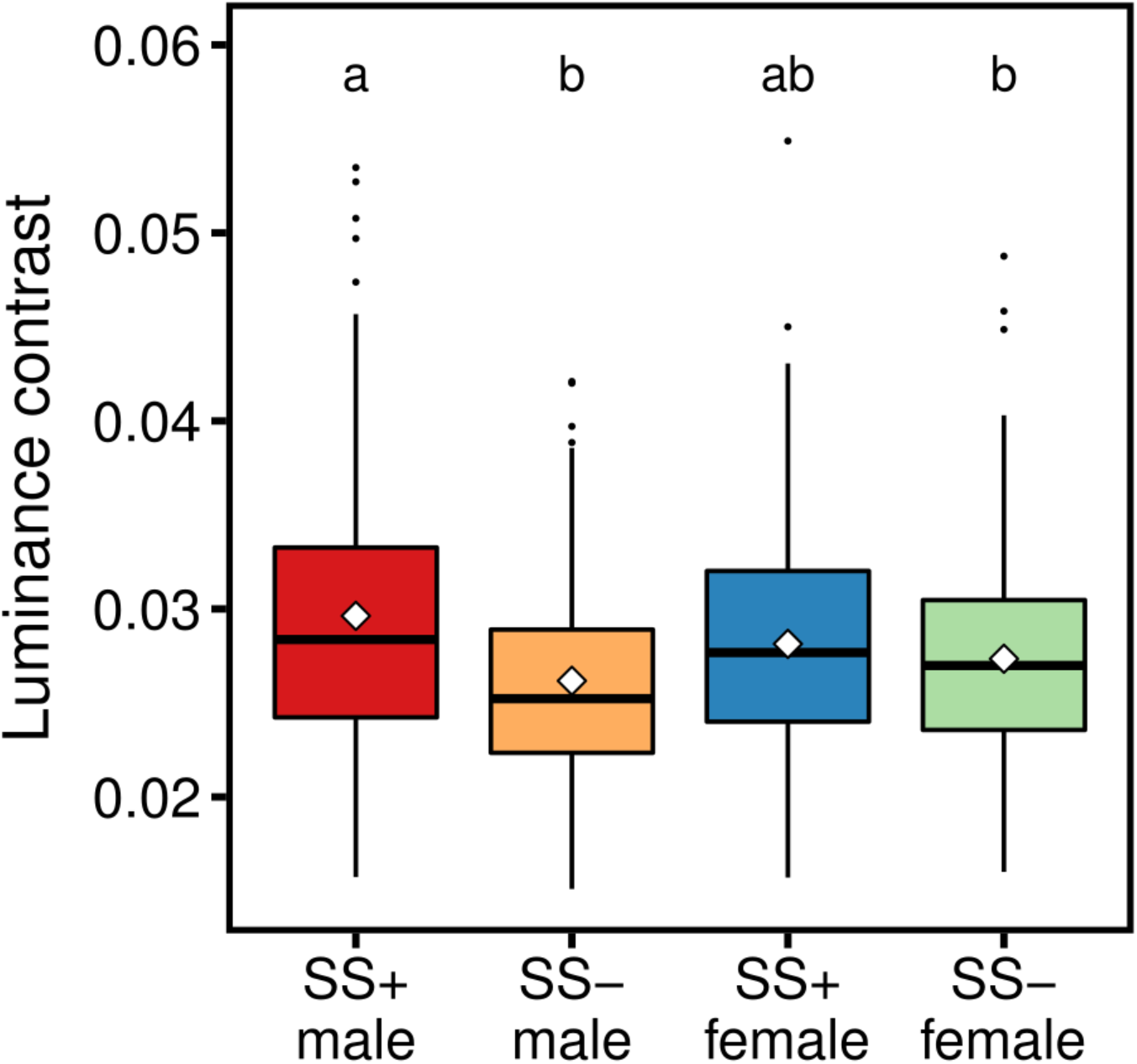
Luminance contrast of WIPs as measured by the standard deviation of the average stimulation the broadband Rh1 photoreceptor in the *Drosophila* visual system. Boxes represent the interquartile range, black bars are medians, white diamonds are means. SS+ = flies from populations evolving with sexual selection, SS- = flies from populations evolving without sexual selection. Differences in letter annotation denote significance at *p*<0.05.

Colour discrimination in *Drosophila* vision is best explained by a system of opponent colour processing, where neurons receive antagonistic input from two or more photoreceptors and the contrast between these inputs is used to process colour information (Schnaitmann *et al.* 2013). To better represent this process we calculated four ‘opponent channels’ that have been empirically validated to accurately describe *Drosophila* colour discrimination (Rh5-Rh3, Rh6-Rh4, Rh6-Rh1, and Rh1-Rh4) (Schnaitmann *et al.* 2013). These were calculated by dividing the cone-catch quanta values of a focal photoreceptor by the sum of the cone-catch quanta values of that photoreceptor and a second comparator photoreceptor (e.g. Rh5/(Rh3+Rh5)) (Kelber *et al.* 2003). We generated images of these opponent channels from cone-catch data and measured the mean hue (average opponent channel pixel values across the wing) and colour contrast (standard deviation in opponent channel pixel values across the entire wing). Due to the high correlation between these four opponent channels (see Methods), we used principal component analyses to extract one significant principal component that explained 90.82% of the variation in the average opponent channel values (i.e. average hue: Table S15), and one significant principal component that explained 79.94% of the variation in colour contrast values (Table S19).

The principal component for average hue described variation in the opponency of long versus short wavelength photoreceptors (Rh5 versus Rh3, and Rh6 versus Rh4), and opponency of narrowband photoreceptors in yellow ommatidia versus broadband photoreceptors (Rh6 against Rh1, and Rh1 against Rh4). The Rh1-Rh4 channel was significantly negatively loaded to this principal component while the remaining 3 channels were significantly positively loaded (Table S15). Thus, higher principal component scores indicate higher reflectance of longer wavelength light (measured by Rh5 and Rh6) relative to shorter wavelength (measured by Rh3 and Rh4), and this relationship becomes stronger as wing luminance (measured by Rh1) increases. Again, we found that the effect of sexual selection on WIPs was different for males and females (GLMM, *χ*^2^_1_=7.51, *p*=0.006). The WIPs of males evolving with sexual selection differ significantly from those of males evolving without (LSMeans, *t* ratio=3.79, p=0.001), showing stronger biases towards longer wavelength light (i.e. visible spectrum), and towards Rh6 and Rh1 in opponency to Rh1 and Rh4, respectively (Figure 4). In contrast, the average hues of female WIPs from either evolution treatment were indistinguishable (LSMeans, *t* ratio=0.06, p=0.99) (Figure 4), did not differ from the no-sexual selection treatment males (LSMeans, *t* ratio<0.29, p>0.99), but were significantly different from males evolving with sexual selection (LSMeans, *t* ratio>-2.71, p<0.04).

**Figure 4.**
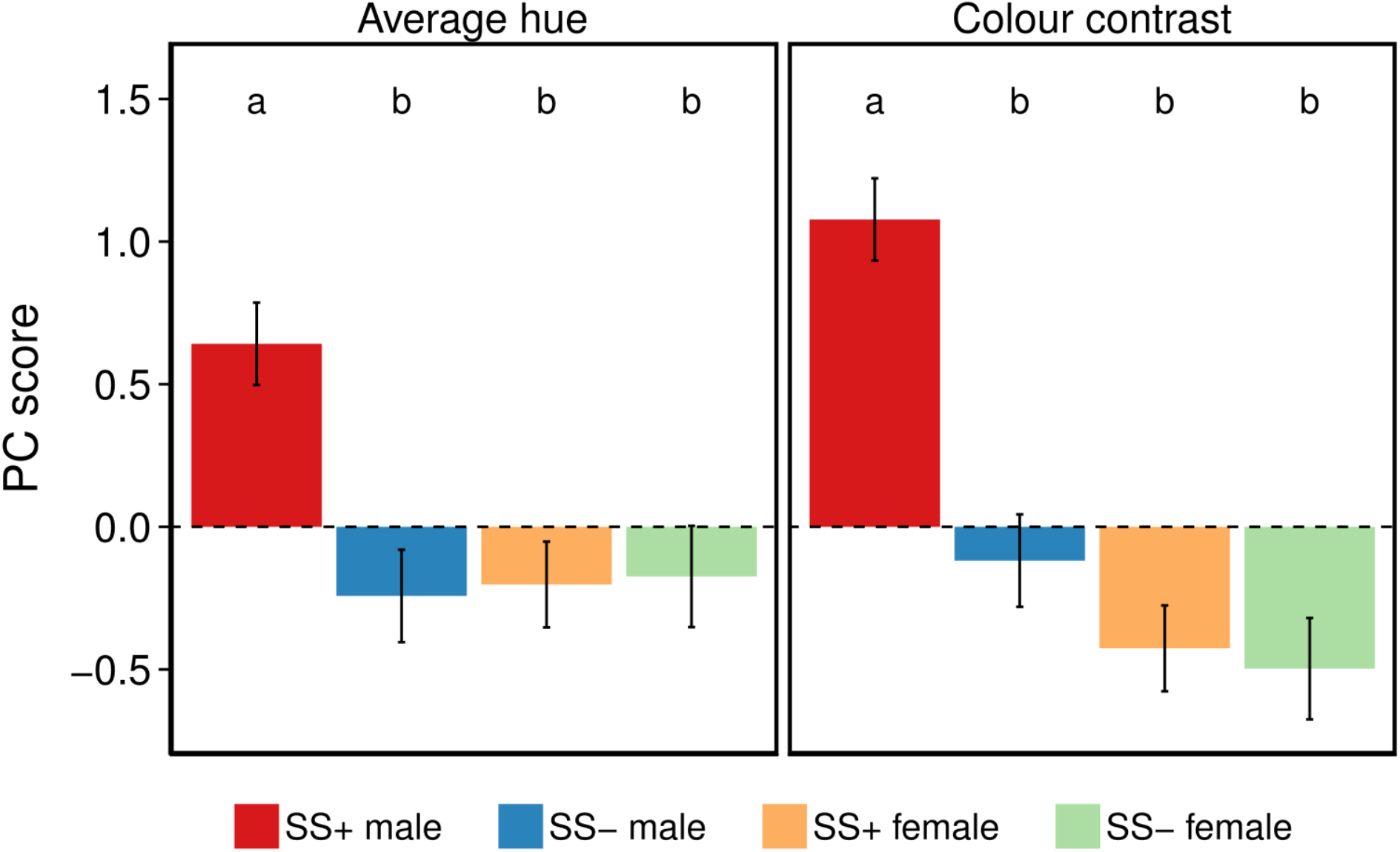
Principal component (PC1) means and standard errors explaining variation in the opponent channels Rh5-Rh3, Rh6-Rh4, Rh6-Rh4, and Rh1-Rh4 for both mean hue (left) and colour contrast (right). SS+ = flies from populations evolving with sexual selection, SS- = flies from populations evolving without sexual selection. Differences in letter annotation denote significance at *p*<0.05.

The principal component for colour contrast describes variation in the same opponent channels as the component for average hue. All four opponent channels were significantly and positively loaded to this principal component (Table S19), and higher principal component scores therefore indicate higher colour contrast levels in all opponent channels. Once again, we found that the effect of sexual selection on WIPs was different for males and females (GLMM, *χ*^2^_1_=19.26, *p*<0.001). The WIPs of males evolving with sexual selection have significantly higher levels of colour contrast than those of males without (LSMeans, *t* ratio=5.42, p<0.001). However, the colour contrast of female WIPs from both selection regimes were indistinguishable (LSMeans, *t* ratio=0.31, p=0.99) (Figure 4), did not differ from the no-sexual selection treatment males (LSMeans, *t* ratio<-1.38, p>0.51), but were significantly different from males evolving with sexual selection (LSMeans, *t* ratio>-5.48, p<0.001).

These results indicate sexual selection has resulted in wing sexual-signal evolution because in all comparisons males from each treatment differed from one another. As such, males from populations that evolved with sexual selection should be more sexually attractive. To test this, the attractiveness (mating latency, a standard measure of male attractiveness: see Methods) of males from experimental populations (when placed with a single virgin tester female) was compared. Ranking population on average attractiveness showed that males from populations evolving with sexual selection (rank sum = 10) were significantly more attractive (better sexual competitors) than males evolving without sexual selection (rank sum = 26) (Mann-Whitney rank sum test: N = 8, Z=-2.31, p=0.02).

## Discussion

Here we show that *D. simulans* wing colour evolved in response to sexual selection when measured using *Drosophila* visual modelling. Critically, this modelling used the full range of human-visible and ultraviolet wavelengths that *Drosophila* can perceive. Behavioural experiments then confirmed that these WIPs correlated with male attractiveness. These results also indicate significant additive genetic variation for WIPs and confirm findings of heritable male attractiveness (Taylor *et al.* 2007). Importantly, the principal component analysis, which effectively summarizes the total data-set into a reduced number of independent variables shows that males evolving with sexual selection have WIPs that are very different from all other flies. While our findings are consistent with those for *D. melanogaster* (Katayama *et al.* 2011) where evidence was found for sexual selection on WIP hue and saturation, that study did not use the explicit model of fly vision or the precise colour measurement we employed here. Studies of sexual colouration in other systems show that failure to consider the appropriate visual system and colour measurement can lead to erroneous conclusions about colour and sexual selection (Bennett *et al.* 1997).

By employing experimental evolution we have explicitly shown that WIPs evolve via sexual selection as males evolving with mate-choice and mate-competition had significantly different wing colouration components than males evolving without sexual selection, and this resulted in males from sexual-selection populations being more attractive to females using a standard measure of attractiveness, latency to mate (Sharma *et al.* 2012a; Sharma *et al.* 2010). Sexual selection resulted in male wings eliciting a stronger response in longer-wavelength light than shorter wavelengths across all four empirically validated opponent channels measured. The wings of males evolving under sexual selection also had higher average luminance (‘perceived brightness’) and luminance contrast than wings from males evolving without sexual selection. Sexual selection therefore seems to favour male wings that have high internal contrast and reflect more light in the human-visible green and blue wavelength regions. However, interpretation of the evolutionary response away from the UV spectrum must be tempered by the low levels of UV light emitted in the controlled environment chambers that housed our populations - this may have constrained evolutionary responses towards the visible spectrum. Despite this, using a standard measure of male attractiveness, males evolving with sexual selection were more attractive to females – they mated faster, and because females determine whether copulation occurs or not (Spieth 1974), mating occurs more rapidly with more attractive males (Taylor *et al.* 2010; Sharma *et al.* 2012a; Taylor *et al.* 2007). It is important to note that we are not implying that WIP evolution is the sole cause of the differences in attractiveness we documented (see e.g. Sharma *et al.* 2012b). However, the covariance between mating-speed and WIP evolution is consistent with WIPs being part of the character-set that in sum defines male sexual attractiveness.

Higher luminance and colour contrast (i.e. variation of WIP luminance and hues) in males evolving with sexual selection can potentially be explained by trade-offs with other sexually and naturally selected phenotypic optima for wing morphology (e.g. flight performance, or acoustic attractiveness in courtship displays) (Radwan 2008; Radwan *et al.* 2016). If selection on wing thickness (which affects WIPs (Shevtsova *et al.* 2011; Katayama *et al.* 2011)) in these other contexts is to some degree orthogonal to selection on WIP colouration from sexual selection, then relaxing sexual selection on WIP colouration could allow these other sources of selection to erode variation in WIP hues that is only relevant in a sexual context. That males from non-sexual selection populations evolve to be more like females - see the principal component analyses - implies that WIPs are costly, which is typical for many sexual traits (Kotiaho 2001). Furthermore, because the mating environment from which our experimental populations were derived includes sexual selection (Taylor *et al.* 2008), the evolution we detect is probably best explained by the relaxing of sexual selection on males in the monogamous populations.

In contrast to males, female wings have the same mean colouration and colour contrast regardless of the selective regime under which they evolved. This is perhaps unsurprising as sexual selection is typically stronger on males (Shuster & Wade 2003; Hosken & House 2011) and our selection protocol only manipulated the opportunity for sexual selection on them. Furthermore, similar sex-specific responses to sexual selection have been found in other *D. simulans* studies (Sharma *et al.* 2012b). While there were some male-female similarities across and within treatments when considering elements of contrast and brightness, the PC analysis that summarizes across all WIPs components showed that males evolving with sexual selection were very different from all other flies.

Taken together, our data suggest that sexual selection drives the evolution of a suite of WIP elements in male *D. simulans.* Specifically, sexual selection favours bright, high contrast, longwave-shifted male WIPs. This finding is further supported by converting raw colour data into empirically validated opponent channels that reflect the neurological processing of colour discrimination in *Drosophila* (Schnaitmann *et al.* 2013). These data suggest that differences between treatments and sexes are an evolutionary response to sexual selection (and its relaxation) on males, and that any intersexual genetic correlation underlying WIPs does not appear to be strong enough to prevent detectably independent sexual evolution (as shown by the PC analysis) although luminance contrast was similar for the sexes within each treatment. Intralocus sexual conflict is a frequent constraint preventing the sexes from reaching sex-specific fitness optima (Rice & Chippindale 2001), but in the *D. simulans* we study, its effects may generally be rather weak (Taylor *et al.* 2010; Sharma *et al.* 2012a), which is consistent with the largely sex-specific WIP responses we document here. A possible exception was seen with luminance contrast, where treatment differences, but not clear sex within-treatment differences, evolved. Nonetheless, the key sex-specific between-treatment comparisons all showed patterns consistent with sexual selection acting most strongly on male WIPs.

## Conclusions

We provide strong evidence for the evolution of WIPs through sexual selection, and we can be reasonably confident that effects are from female mate choice because males from sexual selection lines were more attractive to females. It therefore seems that WIPs are a novel sexual signal that has until very recently have been overlooked in sexual selection research, even in well-studied taxa like *Drosophila.* Our results provide direct evidence that WIPs can evolve in response to sexual selection, and additionally underline the importance of considering the visual sensitivities of intended targets when investigating sexual signals.

## Supporting information

## Acknowledgements

We thank the attendees at the ASAB London Winter Meeting 2017 for comments on these findings. This work was funded by The Leverhulme Trust, BBSRC and NERC, and by the Foundation for Polish Science, International PhD Projects Program co-financed by the European Regional Development Fund within the project MPD/2009-3/5, “Environmental stress, population viability and adaptation”.

## Author contributions

The study was conceived by MDS, JT, NW, JR & DJH. Experiments were conducted by ED, RJ, MFH & JT. Data were analysed by MFH, JT & DJH. The manuscript was written by MFH, JT & DJH. All authors edited and commented on the manuscript.

## Availability of data

Data will be archived in an open source repository upon publication acceptance.

