## Supplementary material for "Sexual selection drives the evolution of wing interference patterns"

**Table S1 Rh1 mean GLMM**

| Effect | $\chi^2$ | df | *p* |
|---|---|---|---|
| Sex | 0.6904 | 1 | 0.406025 |
| Treatment | 6.0885 | 1 | **0.013607** |
| Sex x Treatment | 9.6333 | 1 | **0.001911** |

**Table S2 Rh3 mean GLMM**

| Effect | $\chi^2$ | df | *p* |
|---|---|---|---|
| Sex | 0.0001 | 1 | 0.99178 |
| Treatment | 0.5337 | 1 | 0.46506 |
| Sex x Treatment | 4.5428 | 1 | **0.03306** |

**Table S3 Rh4 mean GLMM**

| Effect | $\chi^2$ | df | *p* |
|---|---|---|---|
| Sex | 0.0108 | 1 | 0.9173 |
| Treatment | 0.9026 | 1 | 0.34209 |
| Sex x Treatment | 5.9194 | 1 | **0.01498** |

**Table S4 Rh5 mean GLMM**

| Effect | $\chi^2$ | df | *p* |
|---|---|---|---|
| Sex | 0.2046 | 1 | 0.650999 |
| Treatment | 5.0333 | 1 | **0.024865** |
| Sex x Treatment | 7.5855 | 1 | **0.005884** |

**Table S5 Rh6 mean GLMM**

| Effect | $\chi^2$ | df | *p* |
|---|---|---|---|
| Sex | 1.4553 | 1 | 0.2276787 |
| Treatment | 7.6007 | 1 | **0.0058347** |
| Sex x Treatment | 12.2213 | 1 | **0.0004725** |

**Table S6 Rh1 SD GLMM**

| Effect | $\chi^2$ | df | *p* |
|---|---|---|---|
| Sex | 1.0496 | 1 | 0.305602 |
| Treatment | 16.3545 | 1 | **0.000052** |
| Sex x Treatment | 6.8375 | 1 | **0.008926** |

**Table S7 Rh3 SD GLMM**

| Effect | $\chi^2$ | df | *p* |
|---|---|---|---|
| Sex | 0.0162 | 1 | 0.8986 |
| Treatment | 0.0397 | 1 | 0.8422 |
| Sex x Treatment | 1.5312 | 1 | 0.2159 |

**Table S8 Rh4 SD GLMM**

| Effect | $\chi^2$ | df | *p* |
|---|---|---|---|
| Sex | 0.0606 | 1 | 0.8055 |
| Treatment | 1.089 | 1 | 0.2967 |
| Sex x Treatment | 1.2811 | 1 | 0.2577 |

**Table S9 Rh5 SD GLMM**

| Effect | $\chi^2$ | df | *p* |
|---|---|---|---|
| Sex | 0.4444 | 1 | 0.50501 |
| Treatment | 16.2752 | 1 | **0.000055** |
| Sex x Treatment | 3.0302 | 1 | 0.08173 |

**Table S10 Rh6 SD GLMM**

| Effect | $\chi^2$ | df | *p* |
|---|---|---|---|
| Sex | 1.4553 | 1 | 0.2276787 |
| Treatment | 7.6007 | 1 | **0.0058347** |
| Sex x Treatment | 12.2213 | 1 | **0.0004725** |

**Table S11 Hue mean LSMEANS**

| | Rh1 | | Rh3 | | Rh4 | | Rh5 | | Rh6 | |
|---|---|---|---|---|---|---|---|---|---|---|
| Treatment/Sex | LSMEAN | SE | LSMEAN | SE | LSMEAN | SE | LSMEAN | SE | LSMEAN | SE |
| SS+ male | 5017.51 | 118.0644 | 4379.27 | 102.0309 | 4802.277 | 96.20561 | 4977.658 | 105.0359 | 5503.996 | 138.0894 |
| SS- male | 4553.283 | 116.7147 | 4203.566 | 101.1849 | 4579.469 | 95.17795 | 4593.687 | 103.7426 | 4887.399 | 136.4793 |
| SS+ female | 4677.583 | 117.1926 | 4246.44 | 101.4873 | 4632.354 | 95.53601 | 4725.194 | 104.2037 | 5014.272 | 137.0408 |
| SS- female | 4729.055 | 116.9221 | 4331.376 | 101.3175 | 4728.714 | 95.33116 | 4763.199 | 103.9438 | 5085.019 | 136.7195 |

**Table S12 Hue mean LSMEANS Tukey adjusted contrasts**

| | Rh1 | | | | | Rh3 | | | | |
|---|---|---|---|---|---|---|---|---|---|---|
| Contrast | Estimate | SE | df | *t* ratio | *p* | Estimate | SE | df | *t* ratio | *p* |
| SS- female - SS+ female | 51.47199 | 117.4639 | 417.3 | 0.438 | 0.9718 | 84.93607 | 86.45572 | 417.66 | 0.982 | 0.7596 |
| SS- female - SS- male | 175.77194 | 127.0491 | 89.46 | 1.383 | 0.5129 | 127.80965 | 91.9375 | 92.06 | 1.39 | 0.5087 |
| SS- female - SS+ male | -288.4555 | 128.3394 | 93.02 | -2.248 | 0.1182 | -47.8944 | 92.90355 | 95.83 | -0.516 | 0.9552 |
| SS+ female - SS- male | 124.29995 | 127.2176 | 90.12 | 0.977 | 0.7628 | 42.87361 | 92.0667 | 92.77 | 0.466 | 0.9664 |
| SS+ female - SS+ male | -339.9275 | 128.5624 | 93.73 | -2.644 | **0.0466** | -132.8305 | 93.07169 | 96.59 | -1.427 | 0.4856 |
| SS- male - SS+ male | -464.2275 | 118.1927 | 420.31 | -3.928 | **0.0006** | -175.7041 | 86.98171 | 420.64 | -2.02 | 0.1822 |

| | Rh4 | | | | | Rh5 | | | | |
|---|---|---|---|---|---|---|---|---|---|---|
| Contrast | Estimate | SE | df | *t* ratio | *p* | Estimate | SE | df | *t* ratio | *p* |
| SS- female - SS+ female | 96.36007 | 92.72983 | 416.91 | 1.039 | 0.7265 | 38.00489 | 108.3237 | 417.45 | 0.351 | 0.9852 |
| SS- female - SS- male | 149.24454 | 103.36142 | 83.18 | 1.444 | 0.4759 | 169.51223 | 115.709 | 92.63 | 1.465 | 0.4627 |
| SS- female - SS+ male | -73.5635 | 104.34635 | 86.33 | -0.705 | 0.8949 | -214.4585 | 116.9163 | 96.39 | -1.834 | 0.2637 |
| SS+ female - SS- male | 52.88447 | 103.48773 | 83.75 | 0.511 | 0.9563 | 131.50735 | 115.8675 | 83.34 | 1.135 | 0.6689 |
| SS+ female - SS+ male | -169.9236 | 104.51559 | 86.95 | -1.626 | 0.3697 | -252.4634 | 117.1252 | 97.15 | -2.155 | 0.1432 |
| SS- male - SS+ male | -222.808 | 93.32013 | 419.77 | -2.388 | 0.081 | -383.9707 | 108.9884 | 420.56 | -3.523 | **0.0027** |

| | Rh6 | | | | |
|---|---|---|---|---|---|
| Contrast | Estimate | SE | df | *t* ratio | *p* |
| SS- female - SS+ female | 70.74693 | 138.9858 | 416.92 | 0.509 | 0.9569 |
| SS- female - SS- male | 197.61956 | 154.6477 | 83.86 | 1.278 | 0.5795 |
| SS- female - SS+ male | -418.9768 | 156.128 | 87.04 | -2.684 | **0.0425** |
| SS+ female - SS- male | 126.87263 | 154.8368 | 84.44 | 0.819 | 0.8451 |
| SS+ female - SS+ male | -489.7238 | 156.3821 | 87.66 | -3.132 | **0.0124** |
| SS- male - SS+ male | -616.5964 | 139.8705 | 419.83 | -4.408 | **0.0001** |

**Table S13 Hue standard deviation LSMEANS**

| | Rh1 | | Rh6 | |
|---|---|---|---|---|
| Treatment/Sex | LSMEAN | SE | LSMEAN | SE |
| SS+ male | 1965.344 | 57.05006 | 2400.291 | 70.87755 |
| SS- male | 1673.539 | 56.27481 | 1959.603 | 69.79065 |
| SS+ female | 1798.53 | 56.54897 | 2096.672 | 70.1686 |
| SS- female | 1735.438 | 56.39345 | 2041.832 | 69.9514 |

**Table S14 Hue standard deviation LSMEANS Tukey adjusted contrasts**

| | Rh1 | | | | | Rh6 | | | | |
|---|---|---|---|---|---|---|---|---|---|---|
| Contrast | Estimate | SE | df | *t* ratio | *p* | Estimate | SE | df | *t* ratio | *p* |
| SS- female - SS+ female | -63.09152 | 61.83114 | 416.98 | -1.02 | 0.7376 | -54.84017 | 81.64773 | 416.63 | -0.672 | 0.9078 |
| SS- female - SS- male | 61.89925 | 66.97815 | 91.06 | 0.924 | 0.782 | 82.22826 | 91.06404 | 86.29 | 0.902 | 0.8033 |
| SS- female - SS+ male | -229.9059 | 67.65762 | 94.68 | -3.398 | **0.0054** | -358.4599 | 91.93628 | 89.53 | -3.899 | **0.0011** |
| SS+ female - SS- male | 124.99077 | 67.06572 | 91.74 | 1.864 | 0.2509 | 137.06843 | 91.17167 | 86.9 | 1.503 | 0.4399 |
| SS+ female - SS+ male | -166.8143 | 67.7745 | 95.4 | -2.461 | 0.0727 | -303.6197 | 92.08409 | 90.17 | -3.297 | **0.0075** |
| SS- male - SS+ male | -291.8051 | 62.21655 | 420.12 | -4.69 | **<0.001** | -440.6881 | 82.17267 | 419.8 | -5.363 | **<0.001** |
